# An in-silico approach for novel molecular glue design by rationalizing known molecular glue mediated ternary complex formation

**DOI:** 10.1101/2022.10.04.510266

**Authors:** Ben Geoffrey A S, Nagaraj M Kulkarni, Deepak Agrawal, Nivedita Bharti, Rajappan Vetrivel, Kishan Gurram

**Affiliations:** Sravathi AI Technology Pvt. Ltd, 63-B, Bommasandra Industrial Area, Bengaluru, Karnataka 560099, India

## Abstract

Protein function modulation using small molecule binding is an important therapeutic strategy for many diseases. However, many proteins remain undruggable due to lack of suitable binding pockets for small molecule binding. Proximity induced protein degradation using molecular glues has recently been identified as in important strategy to target the undruggable proteins. Molecular glues were discovered serendipitously and as such currently lack an established approach for in-silico design rationale. In this work, we attempt to establish the rationale for a known case and having inferred the rationale, we discuss how the rationale can be applied in-silico to design novel molecular glue through AI powered techniques. We believe the establishing of in-silico rationale for molecular glue design would be a valuable and welcome addition to the literature to further accelerate the discovery of molecular glues to drug undruggable targets.

## Introduction

Proteins play important functional roles in the human biological system and in disease their function is either up or down regulated requiring them to be modulated in function using some therapeutic means of intervention. However only 15% of the human proteome remains amenable to modulation of function using small molecules and the rest remain undruggable [1]. An important class of problems which remains challenging or out of scope for the small molecules is the modulation of Protein-Protein Interaction (PPI) using small molecules. This is largely due to the lack of presence of druggable pockets in these proteins [2]. Recently, a certain class of small molecule compounds known as molecular glues were serendipitously discovered and were successfully shown to modulate PPI interaction [3]. However, for rational in-silico molecular glue design there is still no established method(s) reported in the literature [3–6]. In this research we aim to address this gap by appealing to a combination of known and novel in-silico methods for small molecules including NCE generation, NCE optimization, molecular docking, and molecular dynamic simulations for design of molecular glues.

While in-silico approaches for molecular glue design have not been reported in literature, in-silico approach to design hetero-bifunctional molecules such as PROTAC to induce PPI have been reported [7–10]. The hetero-bifunctional molecule must have two functional domains that interact with the two different proteins and bring them together to induce the PPI. While molecular glue type of molecules also conducts a similar function, hetero-bifunctional molecules are usually two ligands connected by a linker, while molecular glues is usually a simple small molecule by construction without a linker but having interacting domains with both the proteins. Therefore, molecular glues enjoy the properties of a small molecule and do not suffer from the draw backs of PROTAC type of molecules which usually have bioavailability and cell permeability issues due to the large size of the molecule. Therefore, it is more desirable to use molecular glues to induce PPI for applications such as protein degradation and hence it is important to have a rational in-silico design approach for molecular glues. We have illustrated an in-silico approach for PROTAC design in our previous work [11] and building on that experience, the design of an in-silico approach for de novo molecular glue design is presented here.

In developing our approach, we rationalize molecular glue mediated ternary complex associated with the PDB ID: 6TD3 using in-silico methods. The molecular glue associated with the ID RC8 recruits DNA damage-binding (DDB) protein 1 to tag and degrade Cyclin dependent Kinase (CDK) which is overexpressed in cancer. The ternary complex formation mediated by RC8 involving the proteins CDK and DDB is investigated in-silico and rationale therein derived has been used to design novel molecular glues for the same problem.

## Methodology

Molecular glues are small molecules that induce two proteins to interact and in the context of ubiquitin ligase this leads to protein degradation and has various therapeutic applications. In the PDB ID: 6TD3, we have a molecular glue that tags Cyclin dependent Kinase (CDK) for ubiquitination and degradation. This leads to the degradation of the cancer-causing protein. For this the small molecule with the RCSB ID: RC8 must induce interaction between DNA damage-binding (DDB) protein 1 and CDK. It does so by forming interactions with the residues of both the proteins in the pocket formed at the interface of both the proteins.

We next describe our in-silico approach step by step.

**Step 1:** The first step in our methodology is to perform protein-protein docking of the two proteins CDK and DDB1 and generate all possible docked poses of the two proteins.

**Step 2:** Next, we perform a throughput scripted pocket analysis to identify whether pockets suitable for small molecules are formed at the interface of all the protein-protein docked poses.

We were able to recover the experimental pose as the leading pose in docking as shown in figure below. DDB1 was taken as the rigid receptor and CDK as the ligand. The experimental pose shown in blue is superimposed on the pose recovered from docking shown in orange in Figure 1 below. The results of the pocket analysis are shown below in Figure 2 where all the pockets formed at the interface of the two proteins suitable for small molecule presence to modulate the PPI is shown in golden color.

**Figure 1:**
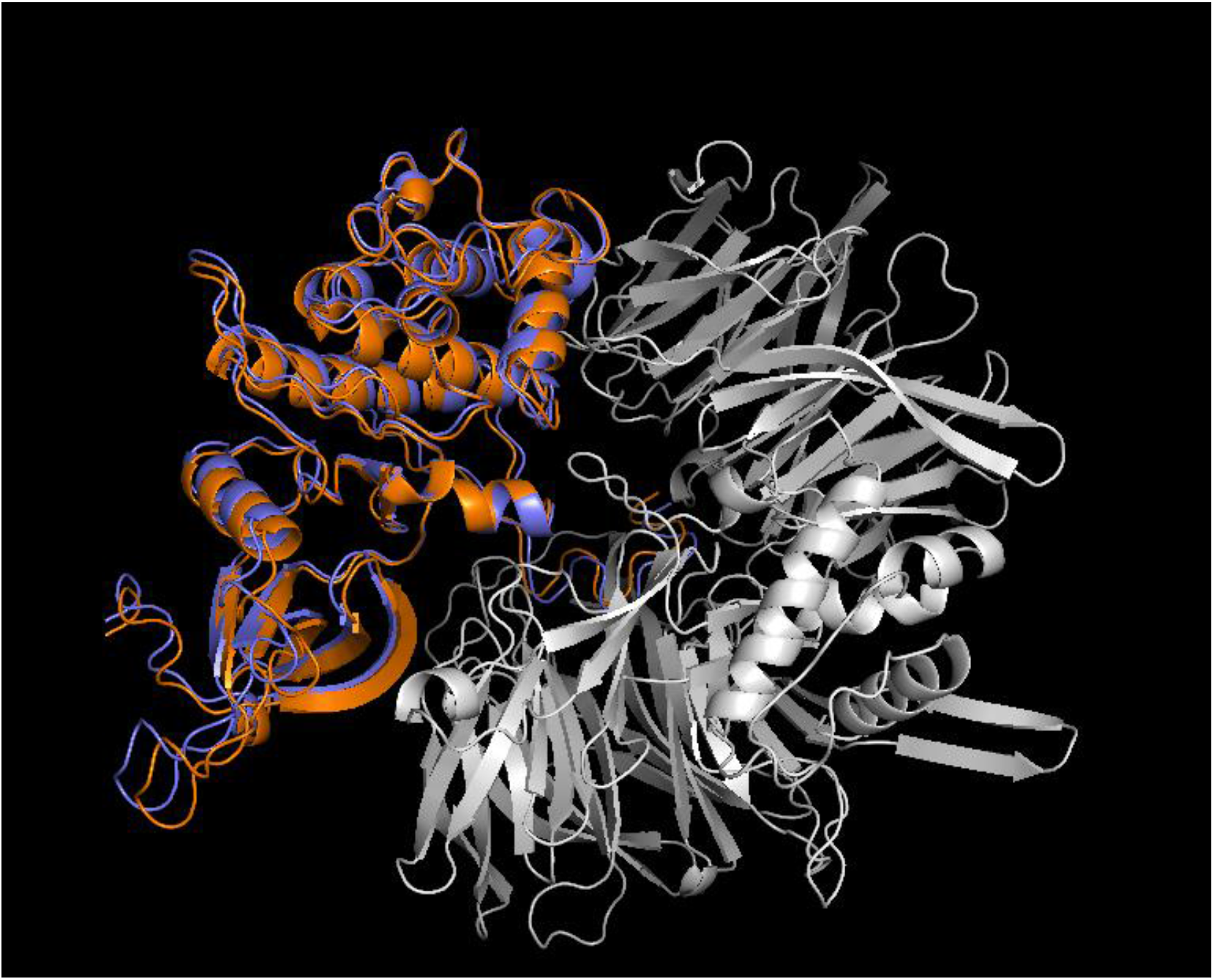
Experimental pose reproduced in protein-protein docking

**Figure 2:**
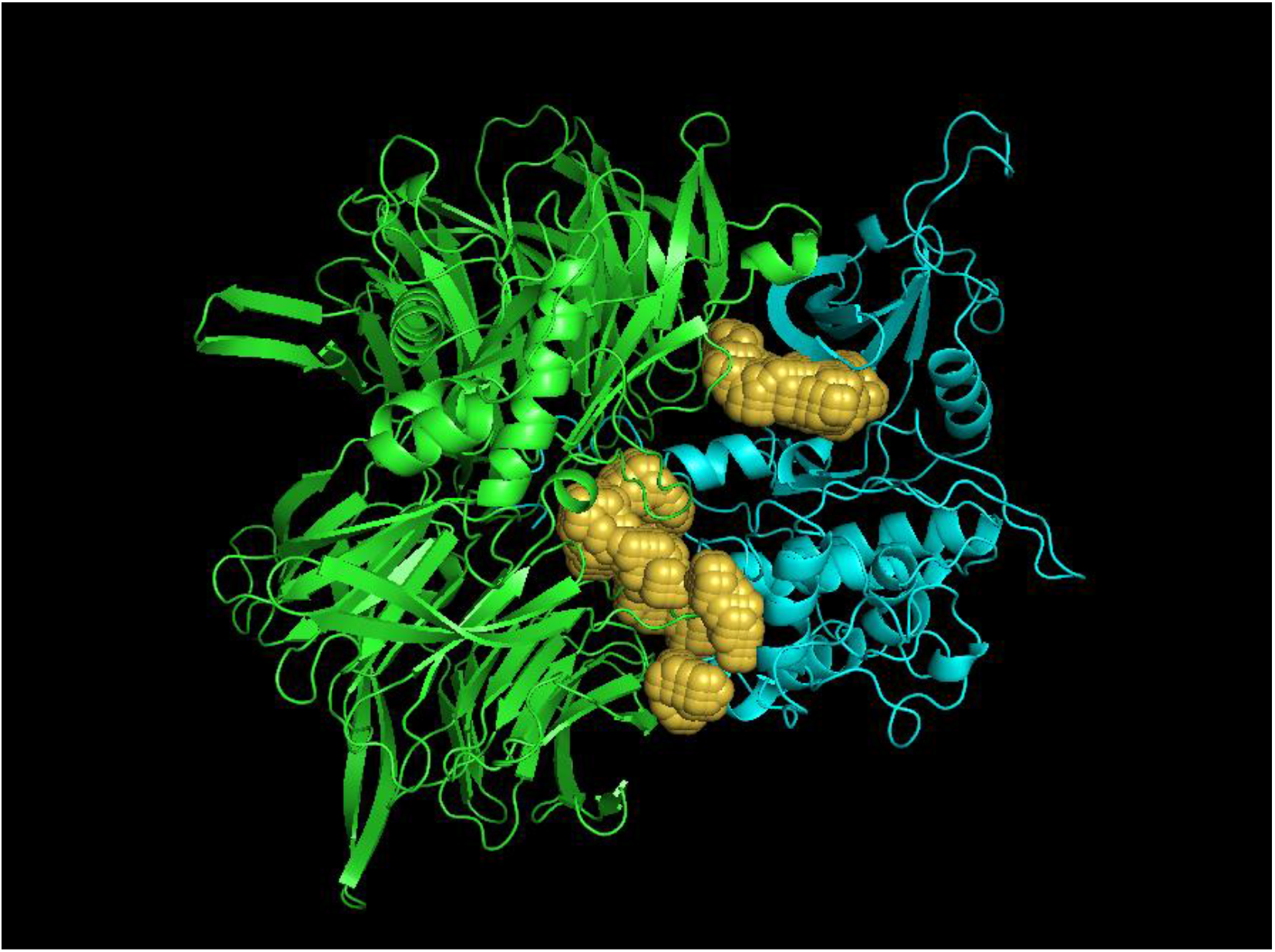
Analysis of the pocket formation at the interface of the two proteins

**Step 3**: Next step in our methodology is to use generative AI to design a small molecule that can bind to the pocket residues of both the proteins with the pocket formed at the interface of both the proteins. We use a mix of generative deep AI to generate small molecules with desirable binding and suitable ADMET properties. This is followed by a docking like method to choose the best among them by generating all conformers of the generated molecules and scoring them with a deep learning-based scoring function for interaction with pocket residues. The deep learning-based scoring functions are trained on datasets such as PDBbind are able to pick up interactions between protein residues and small molecules.

As an illustration of our approach, we perform this process for the known ligand referenced with RCSB ID: RC8 and obtain the experimental pose with all interactions intact and is shown in Figure 3 below. The ligand has interactions with residues from protein residues belonging to chain A and B of the two proteins.

**Figure 3.**
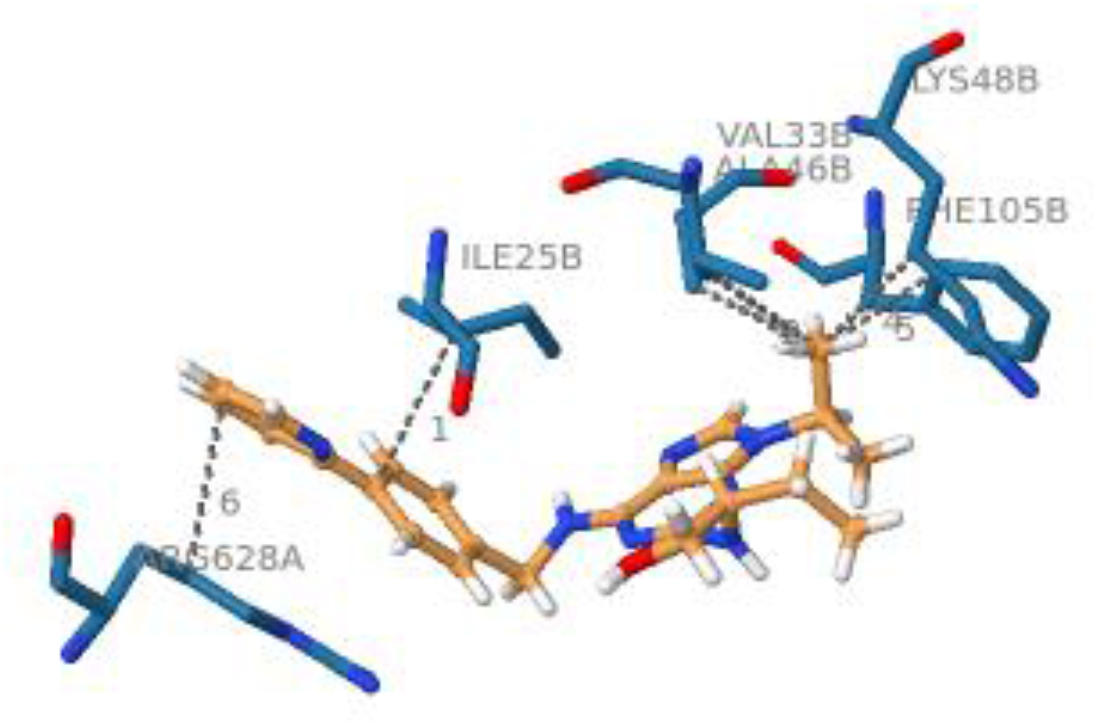
Molecular glue mediated interactions

**Step 4**: To further analyze in-silico, whether the small molecule is able to induce and stabilize the PPI we perform a long Molecular dynamics (MD) simulation and check whether the small molecule interactions with the pocket residues are intact and whether the ternary complex is stable. We found that the known ligand referenced with RCSB ID: RC8 aided in stable ternary complex formation and the entire system was stable during MD. Further, in-silico results of the molecular glues designed by us using our design approach involving small molecules suited to bind to the residues of interest and with desirable ADMET properties are given below in results and discussion section. Our entire methodology is captured as a workflow in the diagrammatic flow in Figure 4 below.

**Figure 4.**
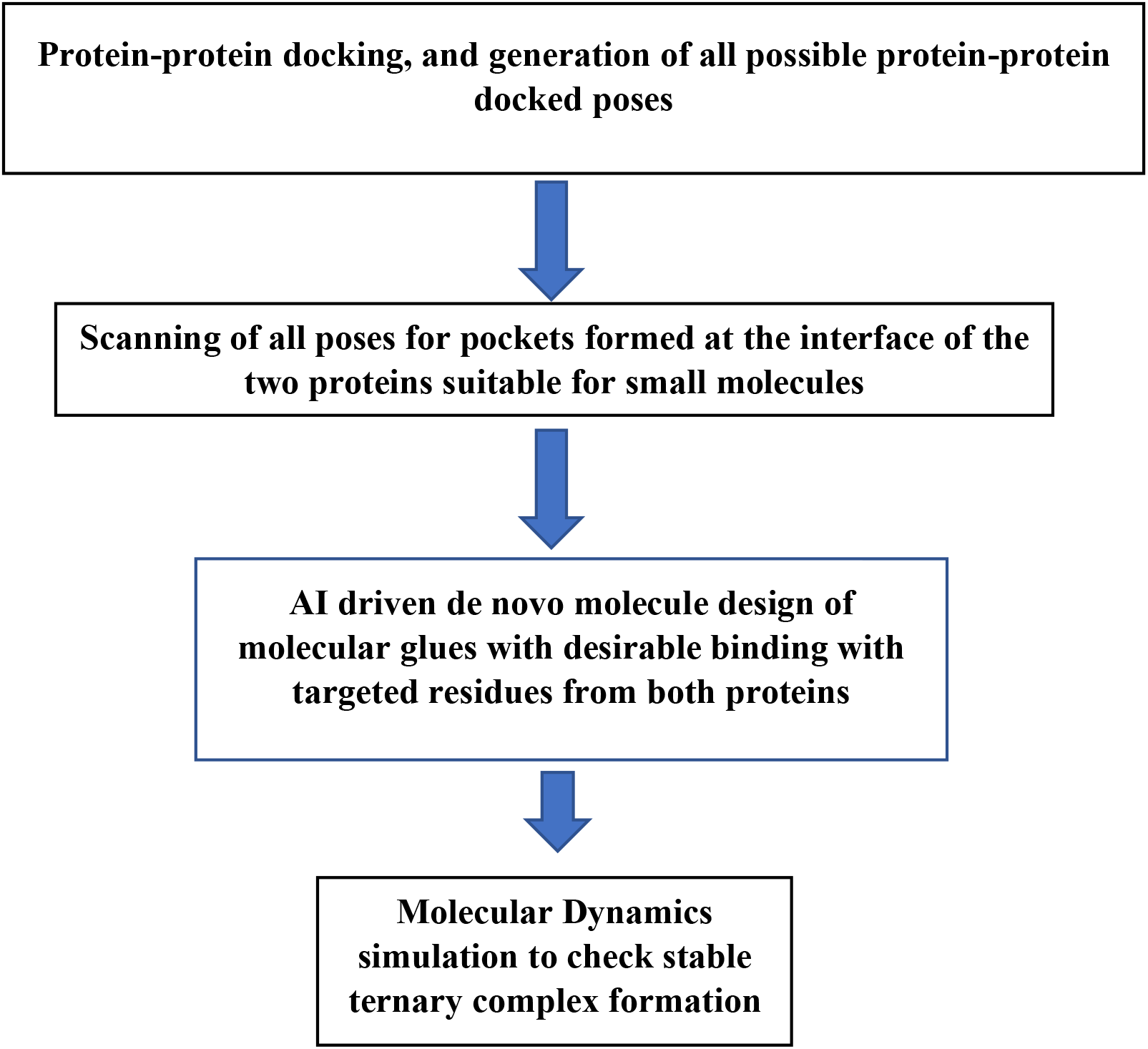
Molecular glue design workflow

## Results and discussion

In order to rationalize the formation of RC8 mediated ternary complex formation involving CDK and DDB, we performed a molecular dynamics simulation and analyzed the ability of the RC8 to mediate stable ternary complex formation of CDK and DDB and found that the ternary complex mediated by RC8 was stable in 50 ns long MD simulation. The RMSD stabilization plot is shown below in Figure 5. The fluctuations in the RMSD associated with the structure of the ternary complex become minimal over time meaning there is minimal change in the position of the atoms and therefore the structure of the ternary complex is stable. The small molecule RC8 was able to mediate this stable ternary complex formation and it is further rationalized through analysis of its interaction with the residues from both the proteins at the protein-protein interface. Though these two proteins natively are energetically not favored to interact, the molecular glue by interacting with residues from both proteins at the protein-protein interface, induce energetics in favor of interaction and ternary complex formation. This is displayed below in Figure 5.

**Figure 5.**
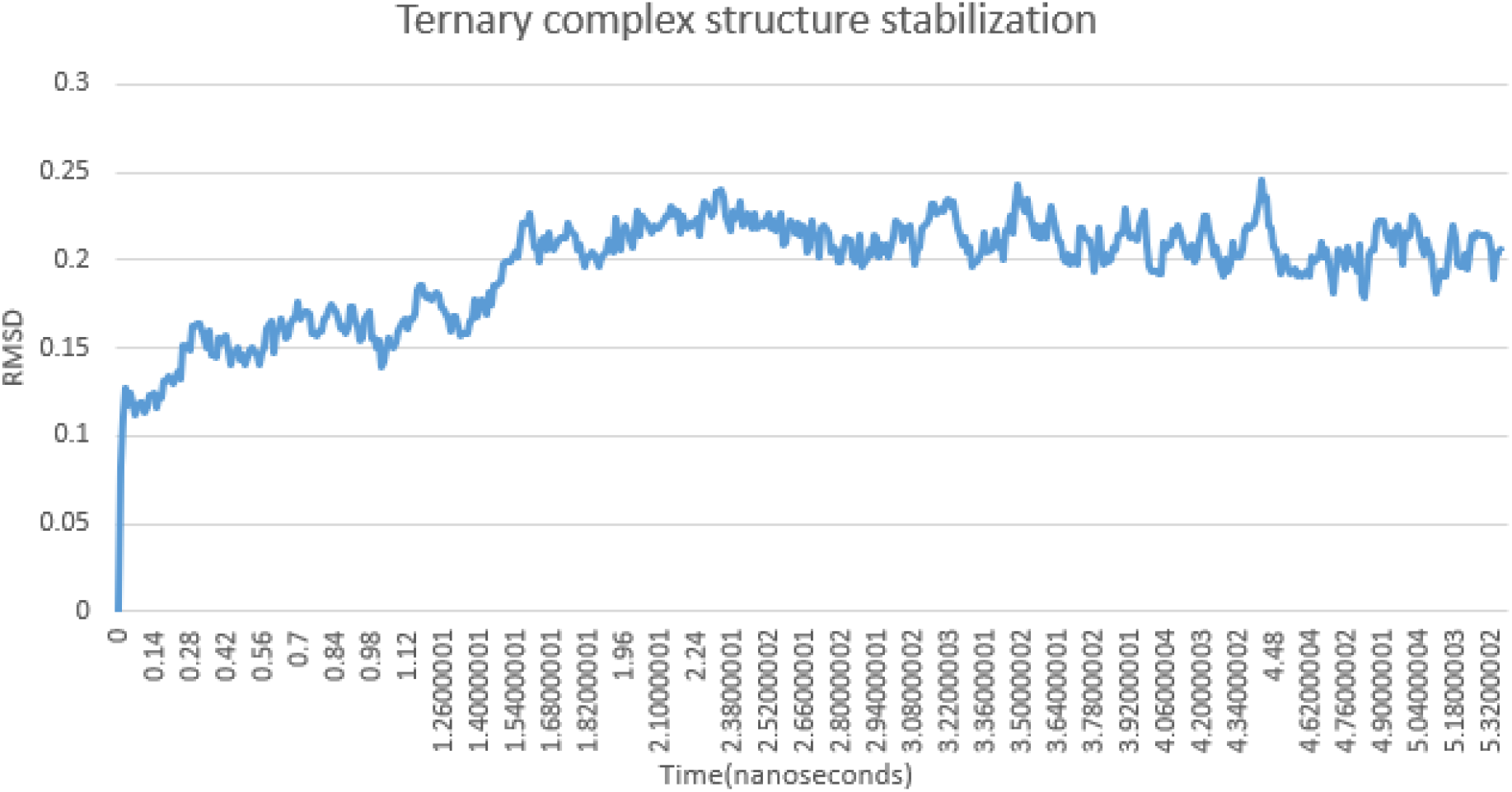
Structure stabilization of ternary complex

The ability of RC8 to mediate interaction between CDK and DDB is rationalized to be due to RC8 having interactions with residues of CDK and DDB which is depicted in the visualization shown below in Figure 6.

**Figure 6.**
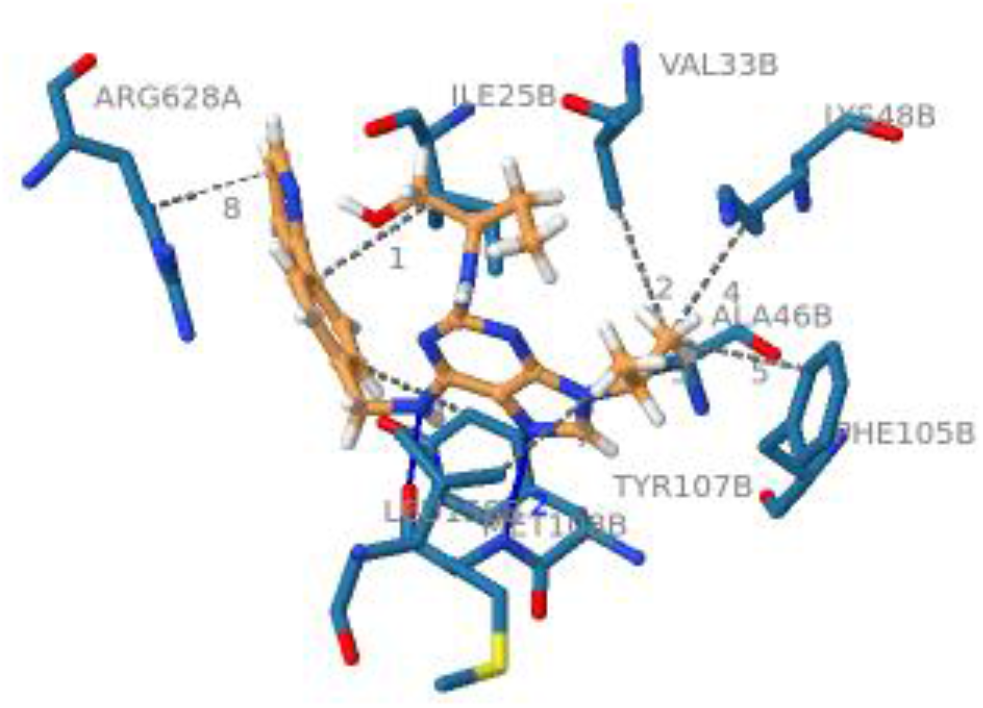
RC8 interacting with DDB and CDK

In Table 1 below the interacting residues are tabulated.

**Table 1.**
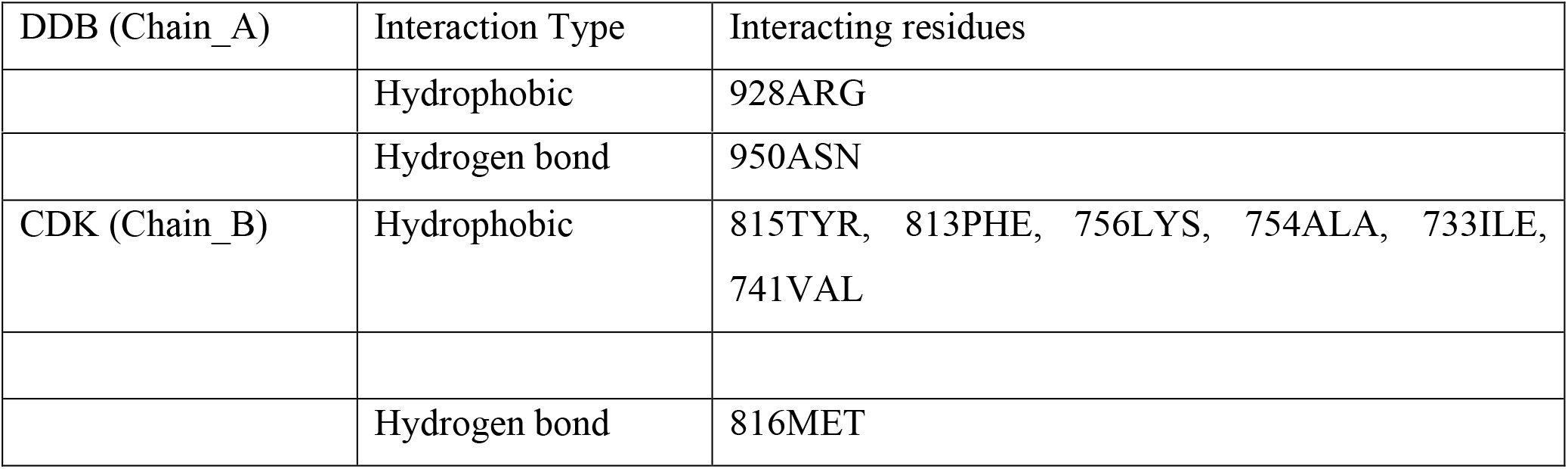
RC8 interaction with DDB and CDK

Interaction of RC8 with CDK is strong due the presence of druggable pocket and though the molecular glue weakly interacting with DDB, it is sufficient to induce proximity of DDB with CDK to initiate degradation. The ability of molecular glues to act via proximity-based mechanism opens the scope of using molecular glues to drug undruggable targets via degradation initiated by molecular glue mediated close proximity of E3 ligase and target of interest.

Having validated our in-silico rationalization of molecular glue induced PPI through the known example associated with PDB ID: 6TD3, we next used our AI driven de novo molecular design approach to design molecule glues with desired binding properties and favorable ADMET properties following the protocol mentioned in STEP 3. However, since drug repurposing is quicker approach to drug discovery, we screened the potential of purchasable drugs to be able to modulate the PPI interaction and mediate the formation of the ternary complex. The hit compounds were modified and optimized using deep generative AI models to give potentially patentable molecular glues to modulate the PPI interaction involving CDK and DDB and initiate the degradation of CDK. The results of the leading molecular glue candidates to induce interaction between CDK and DDB1 are shown below in Table 2

**Table 2.**
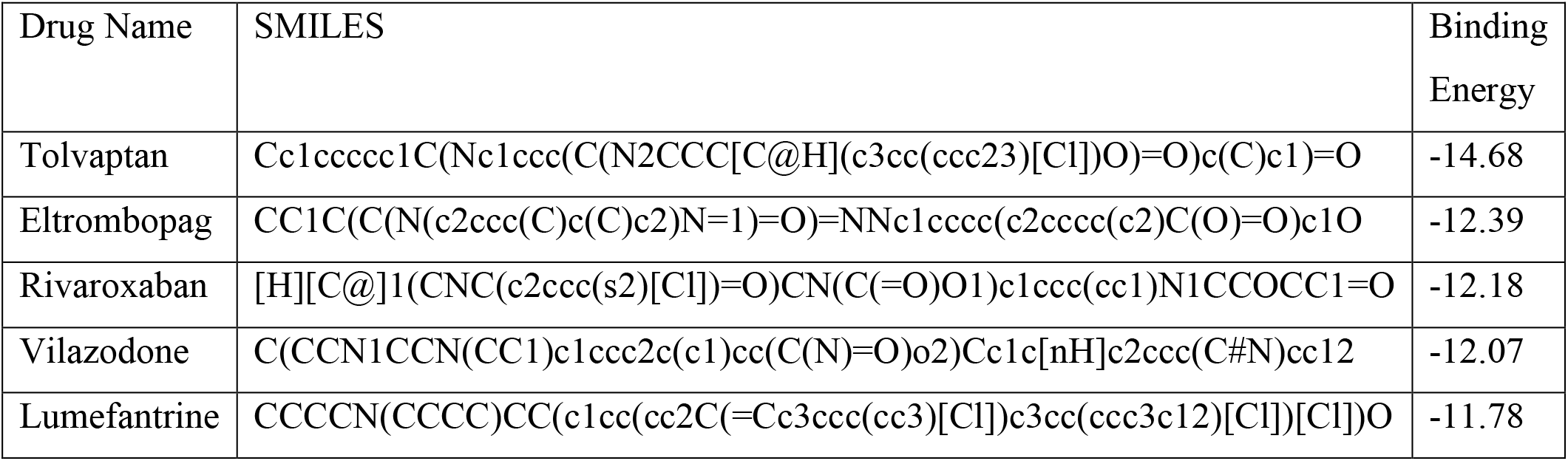
Potential Molecular glue candidates for CDK degradation among purchasable drugs

In Figure 7 we show the ability of a potential molecular glue candidate Tolvaptan to induce PPI via its ability to interact with residues of both the proteins at the pocket formed at their interface.

**Figure 7.**
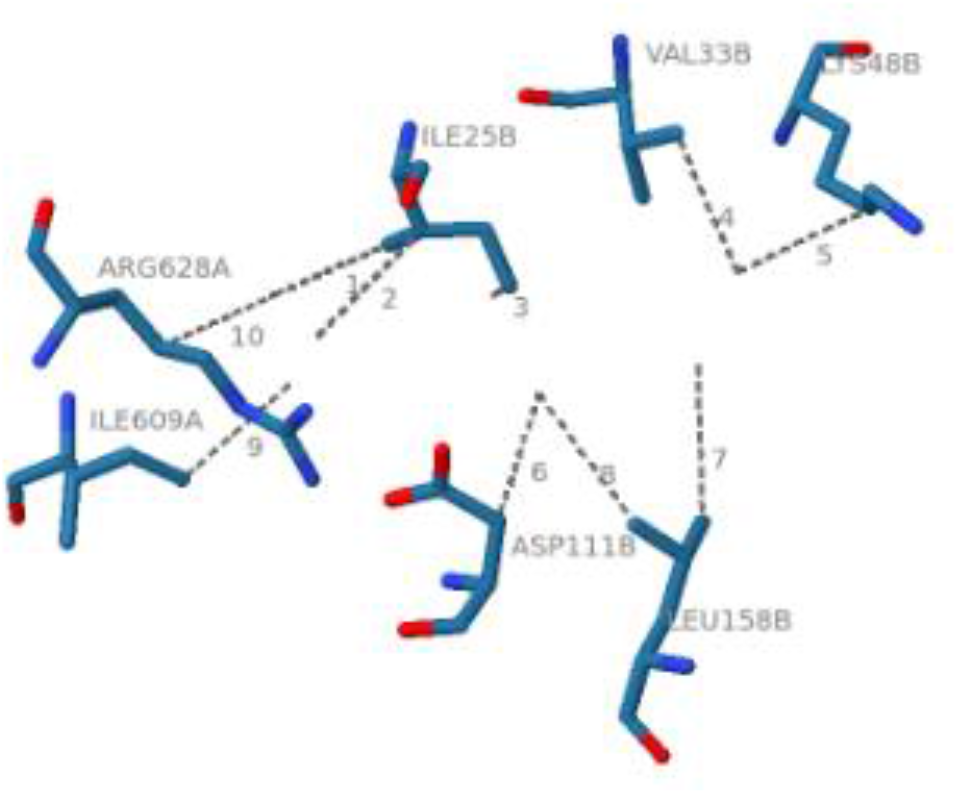
Tolvaptan interacting with DDB and CDK

The drug Tolvaptan interacts with the residues of both CDK and DDB which are tabulated in Table 3 below.

**Table 3.**
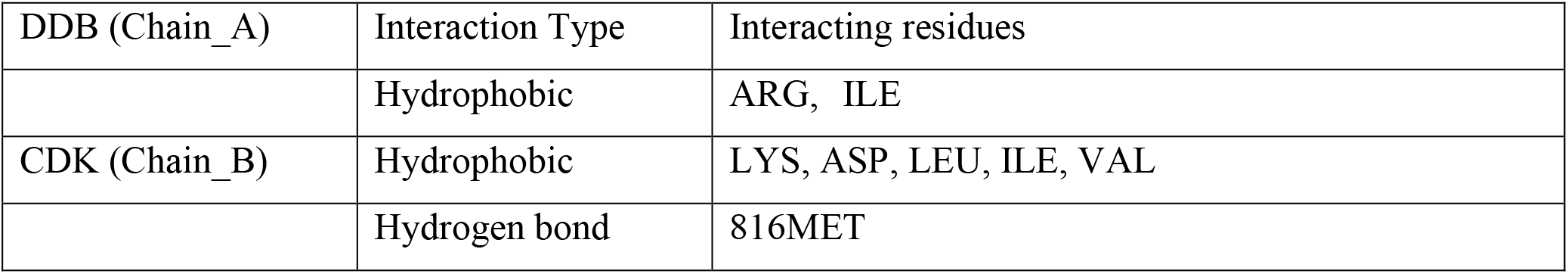
Tolvaptan interactions with DDB and CDK

Also, we further validated the ability of Tolvaptan to mediate the stable ternary complex formation involving CDK and DDB. We found that Tolvaptan mediated a stable ternary complex involving CDK and DDB in a 50-nanosecond long MD simulation and the RMSD stabilization of Tolvaptan along with the reference RC8 is shown in Figure 8 below.

**Figure 8.**
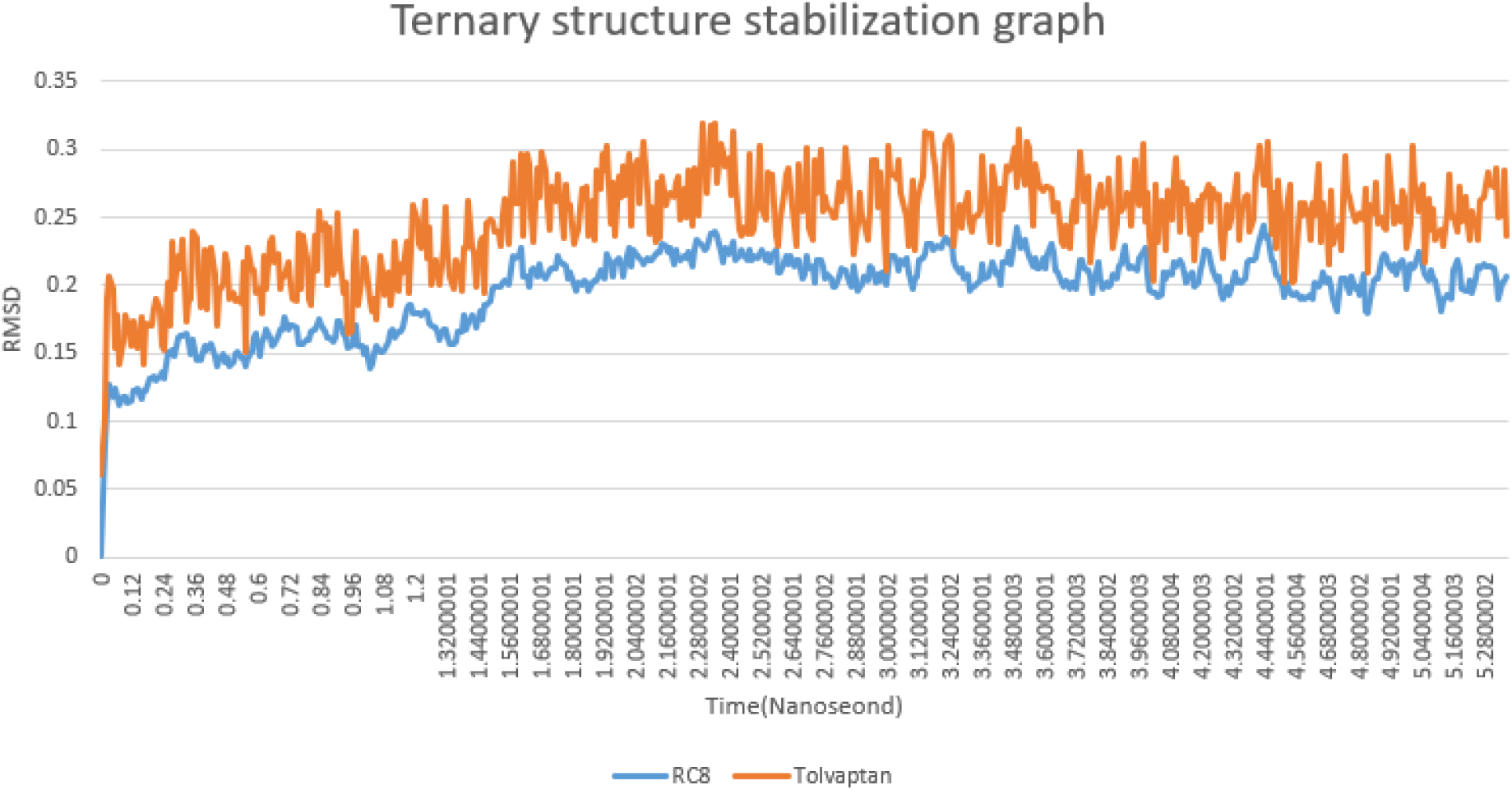
Ternary complex stabilization plot

Furthermore, building on the known approved drugs, we used our deep generative AI approach to generate more promising NCE molecular glues for CDK degradation which are tabulated in Table 4 below.

**Table 4.**
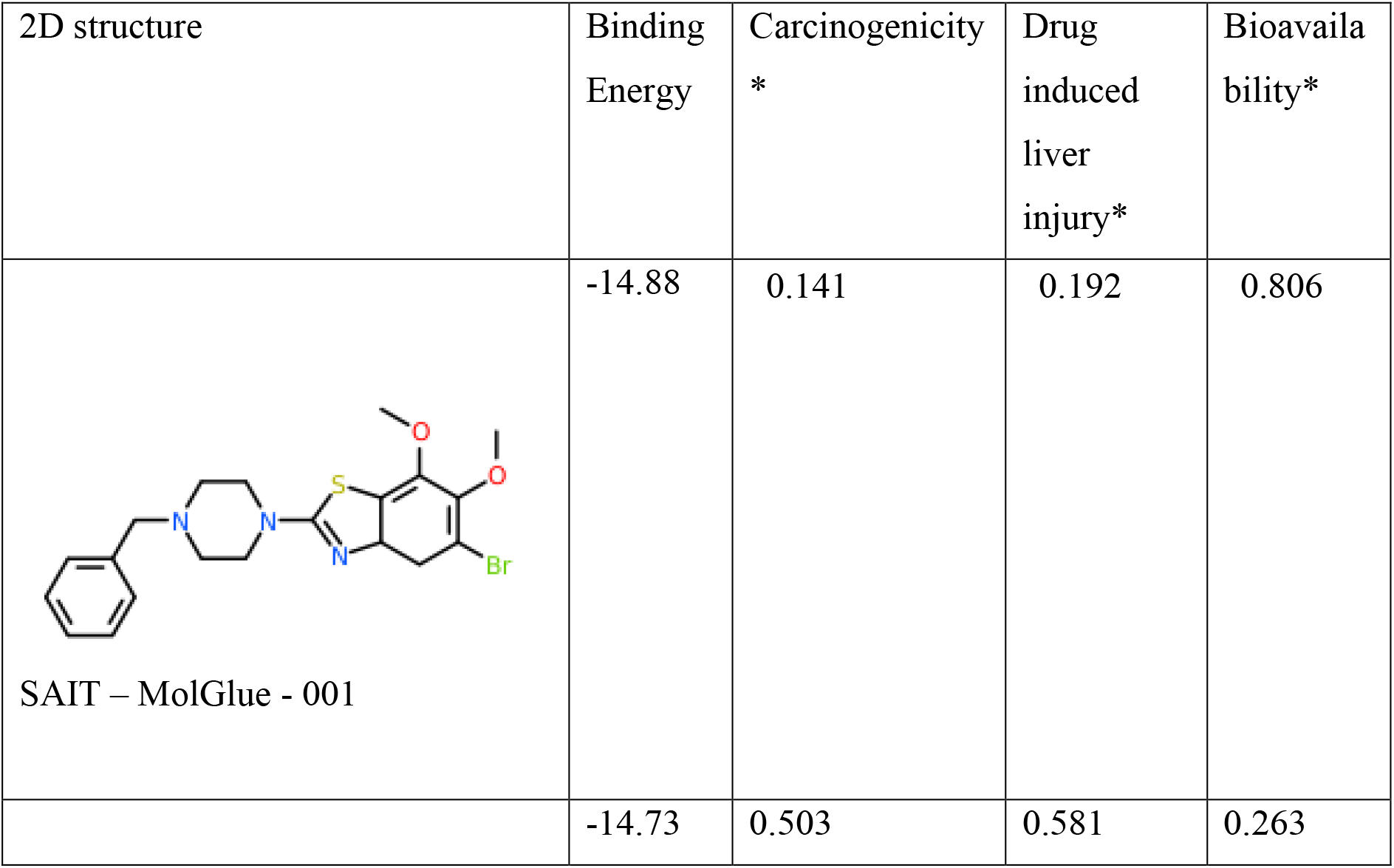
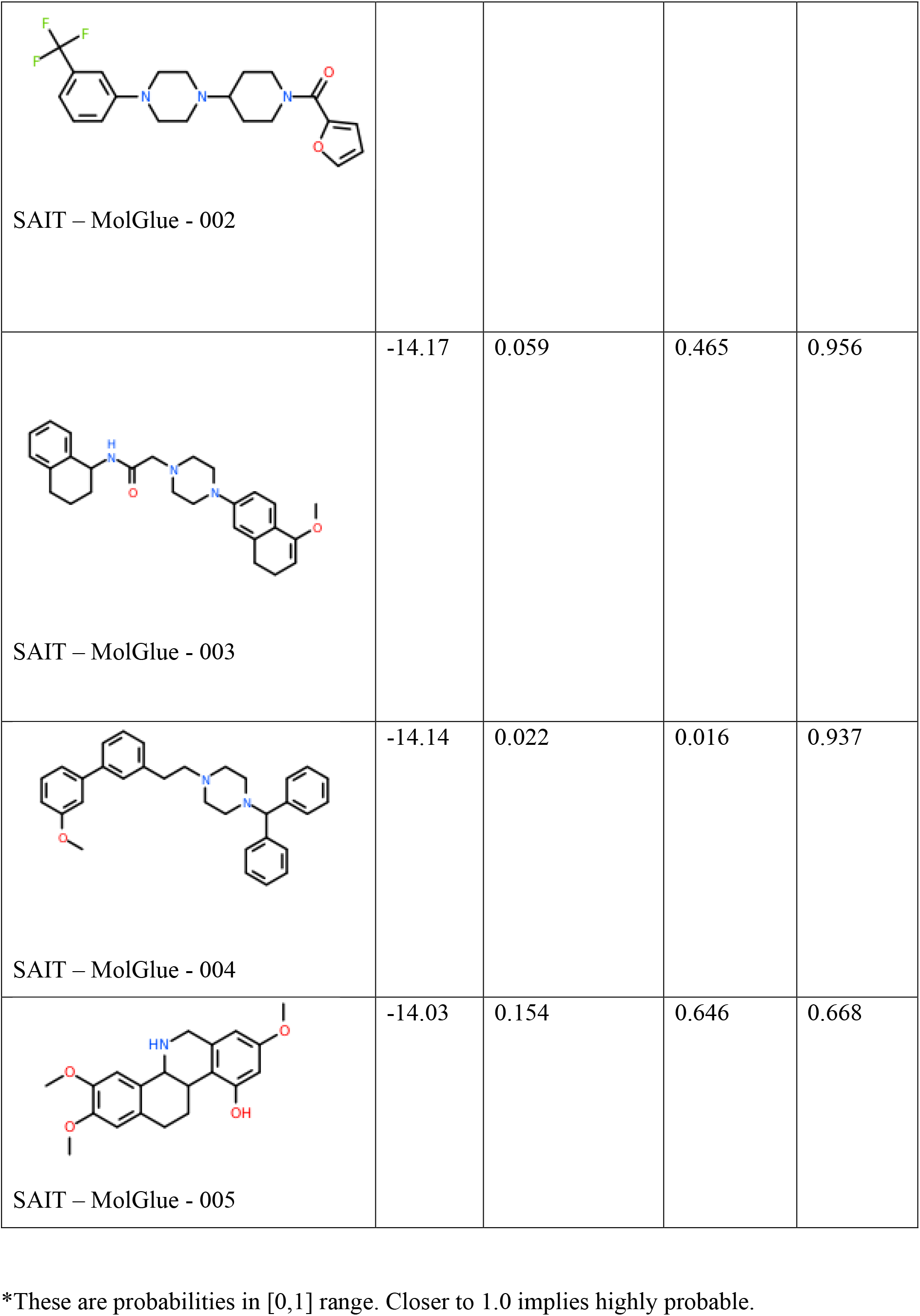
Potential NCE Molecular glue candidates for CDK degradation

As per our in-silico rationale, an ideal molecular glue candidate should pose binding domains with both the proteins and thereby modulate the PPI/proximity between the protein of interest and E3 ligase to initiate degradation. Therefore, in the generation of NCEs through Generative AI we have optimized for NCEs that have strong non-covalent interactions between protein of interest and the E3 ligase. The properties of these generated NCEs were also optimized for less toxicity and better bioavailability. Generated candidates with their in-silico binding, toxicity and bioavailability predictions are tabulated in Table 4. The binding energy contribution comes from interaction with residues of both the proteins at the pocket formed at the interface of both the proteins.

For the first candidate listed in the Table 4, a detailed breakup of the interactions is given below in Table 5. It is observed that the hydrogen bonds are formed with residues from both the proteins which would contribute to the stability of the PPI. Further the other non-covalent interactions picked up Hydrophobic, Salt Bridges and Halogen bond would also contribute to induce the proximity induced degradation cascade.

**Table 5.**
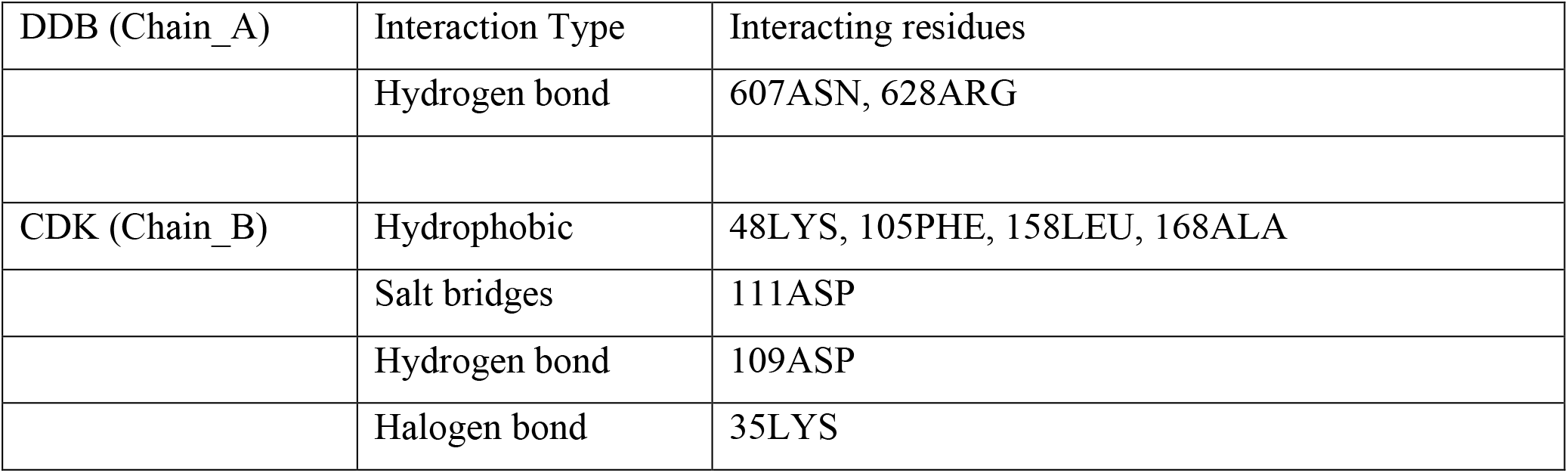
Tolvaptan interactions with DDB and CDK

## Conclusion

Molecular glues which are closer in resemblance to traditional small molecules are a very lucrative alternative to heterobifunctional PROTACs for target protein degradation as PROTAC space remains plagued by the permeability and the bio-availability problem which arise due to the large sizes of PROTAC molecules. However, for molecular glues that were discovered serendipitously a rational design approach remains still to be established. To establish the in-silico rationale for molecular glue design, we take a known molecular glue associated with the PDB ID: 6TD3 and establish the in-silico design rationale for the already discovered molecular glue RC8. Further, we use an AI driven de novo molecular design toolkit to design molecular glues with desired in-silico binding and ADMET properties. This establishing of in-silico approach to molecular glue design will accelerate high-throughput screening and designing of molecular glues. Further open a new avenue for the application of in-silico design principles and methodologies.

Although we did not attempt free energy calculations associated with the molecular glue mediated ternary complexation event here, such work would add quantitative rationale decision making value in molecular glue design, which we hope to conduct in our subsequent research work.

No work is without shortcomings, especially first few attempts at a problem and to our knowledge this is the first work to establish an in-silico rationale, throughput design and screening approach for molecular glues and therefore we believe any unreached goals in this work would be reached by subsequent work from our group or others in the community. More research for establishing in-silico methods for molecular glue design along the lines we have attempted here are required to improve our understanding and further validate the in-silico methods for drug design of molecular glues.

## Acknowledgements

We are grateful for the various useful discussions we have had with our colleagues at Sravathi Artificial Intelligence. In particular, we would like to thank Srinivasan Krishnaswami, Sourabh Mundra and Kondabala Rajesh for their valuable input and insights.

## Conflict of Interest disclosure

All authors are employees of Sravathi AI Private Limited, Bengaluru, India. We do not have any conflicts of interest to report.

## Notes

### Competing Interest Statement

The authors have declared no competing interest.

## References

1. Kana, Omar, and Michal Brylinski. “Elucidating the druggability of the human proteome with eFindSite.” Journal of computer-aided molecular design 33.5 (2019): 509–519.

2. Qiu, Yuran, et al. “Computational methods-guided design of modulators targeting protein-protein interactions (PPIs).” European Journal of Medicinal Chemistry 207 (2020): 112764.

3. Domostegui, Ana, et al. “Chasing molecular glue degraders: screening approaches.” Chemical Society Reviews (2022).

4. Sasso, Janet M., et al. “Molecular glues: The adhesive connecting targeted protein degradation to the clinic.” Biochemistry (2022).

5. Geiger, Thomas M., et al. “Clues to molecular glues.” Current Research in Chemical Biology 2 (2022): 100018.

6. Maneiro, M., et al. “PROTACs, molecular glues and bifunctionals from bench to bedside: Unlocking the clinical potential of catalytic drugs.” Progress in medicinal chemistry 60 (2021): 67–190.

7. Drummond, Michael L., and Christopher I. Williams. “In silico modeling of PROTAC-mediated ternary complexes: validation and application.” Journal of chemical information and modeling 59.4 (2019): 1634–1644.

8. Weng, Gaoqi, et al. “Integrative modeling of PROTAC-mediated ternary complexes.” Journal of Medicinal Chemistry 64.21 (2021): 16271–16281.

9. Zaidman, Daniel, Jaime Prilusky, and Nir London. “PRosettaC: Rosetta based modeling of PROTAC mediated ternary complexes.” Journal of chemical information and modeling 60.10 (2020): 4894–4903.

10. Liao, Junzhuo, et al. “In Silico Modeling and Scoring of PROTAC-Mediated Ternary Complex Poses.” Journal of Medicinal Chemistry 65.8 (2022): 6116–6132.

11. AS, Ben Geoffrey, et al. “A New In-Silico Approach for PROTAC Design and Quantitative Rationalization of PROTAC mediated Ternary Complex Formation.” bioRxiv (2022).

