## Supplemental for "An in-silico approach for novel molecular glue design by rationalizing known molecular glue mediated ternary complex formation"

**Supplementary with 8 Figures and 5 Tables**

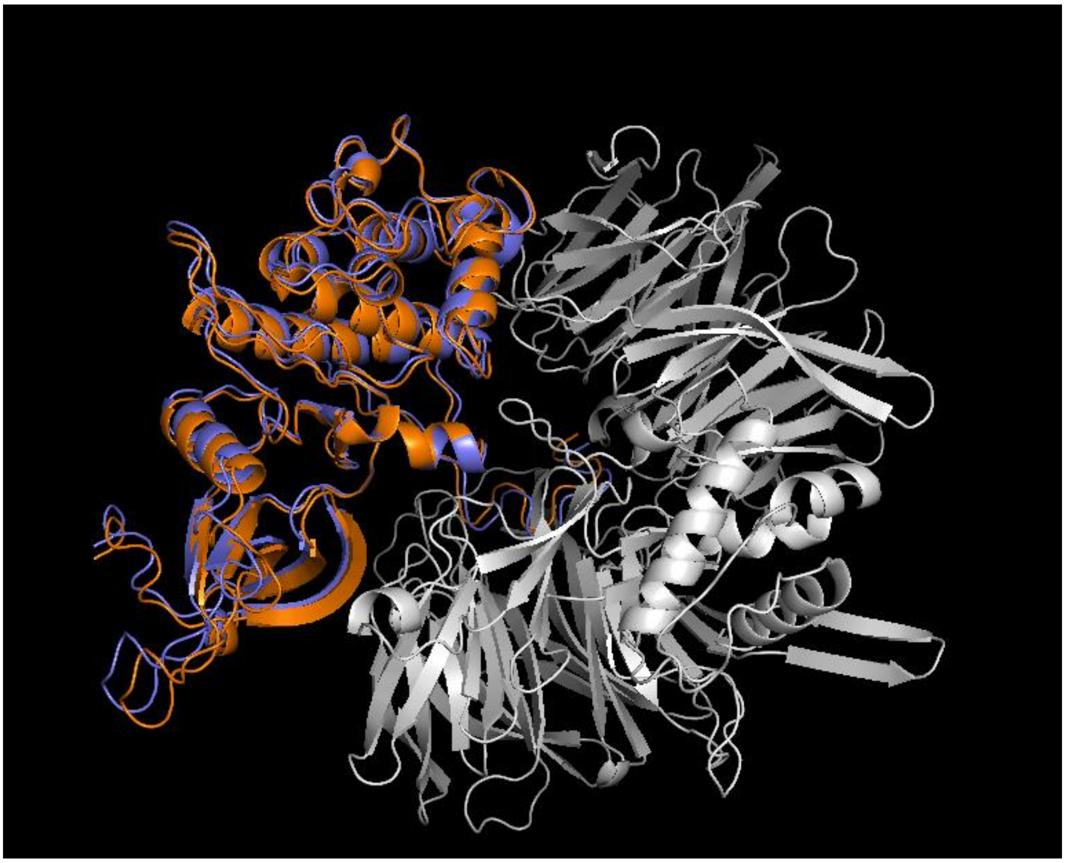

Figure 1: Experimental pose reproduced in protein-protein docking

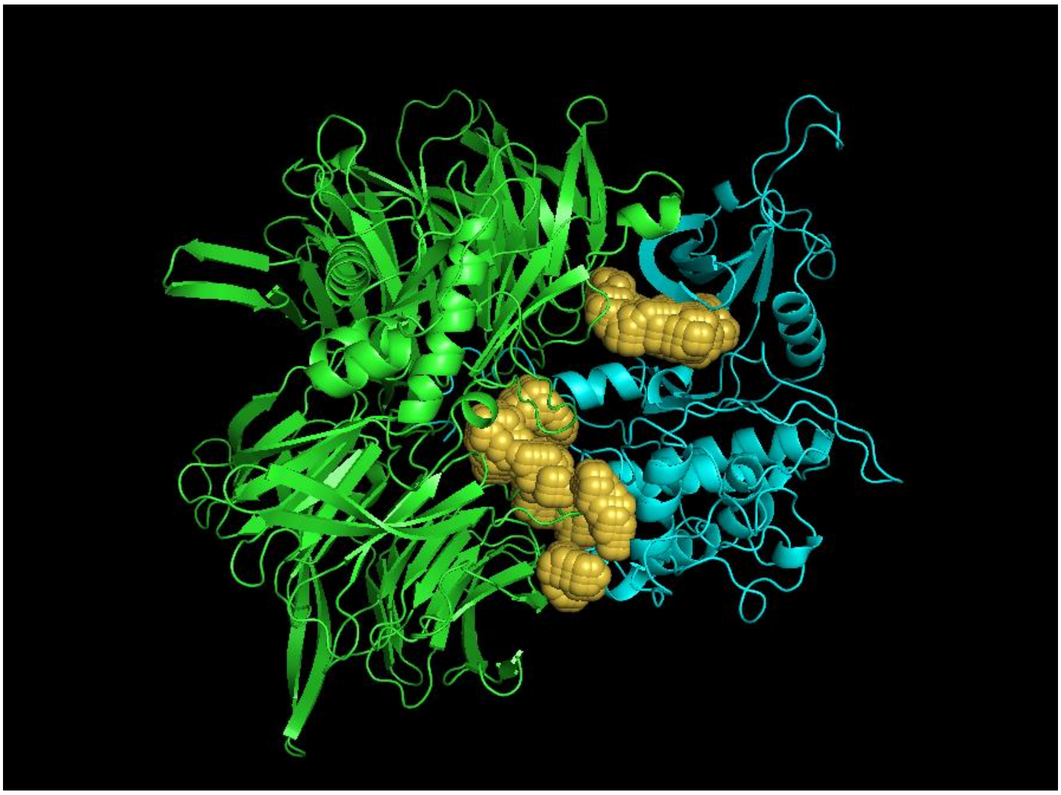

Figure 2: Analysis of the pocket formation at the interface of the two proteins

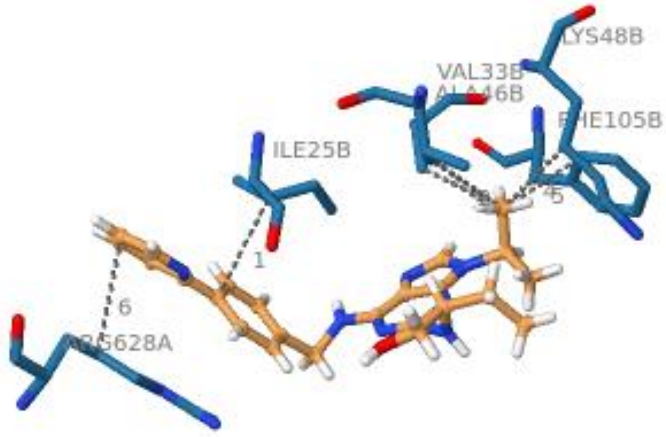

Figure 3 – Molecular glue mediated interactions

**Protein-protein docking, and generation of all possible protein-protein docked poses**

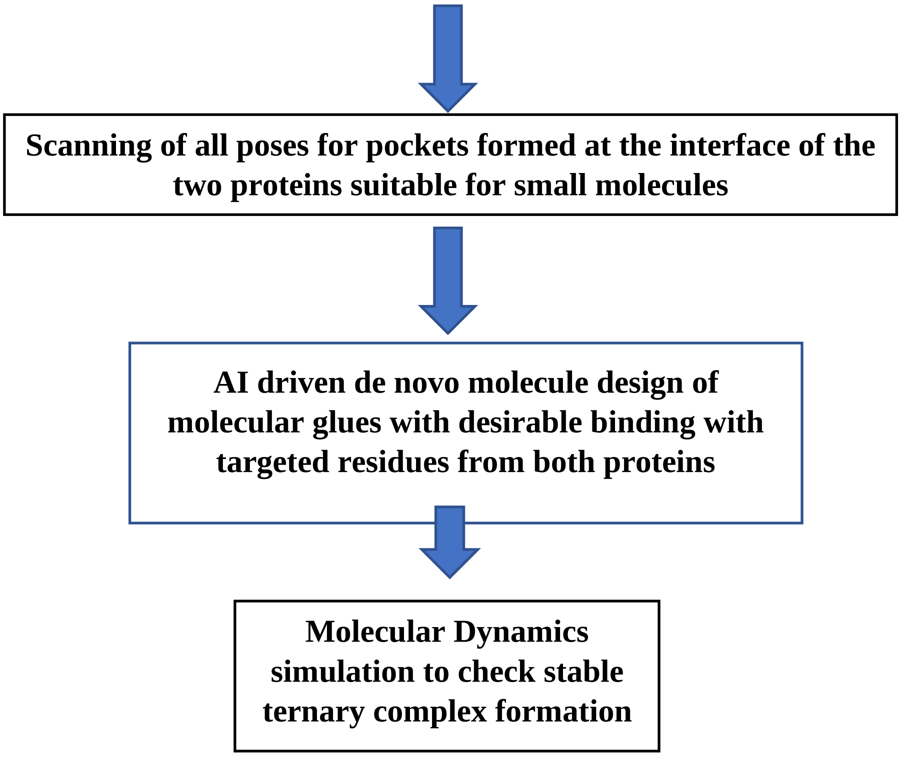

Figure 4 – Molecular glue design workflow

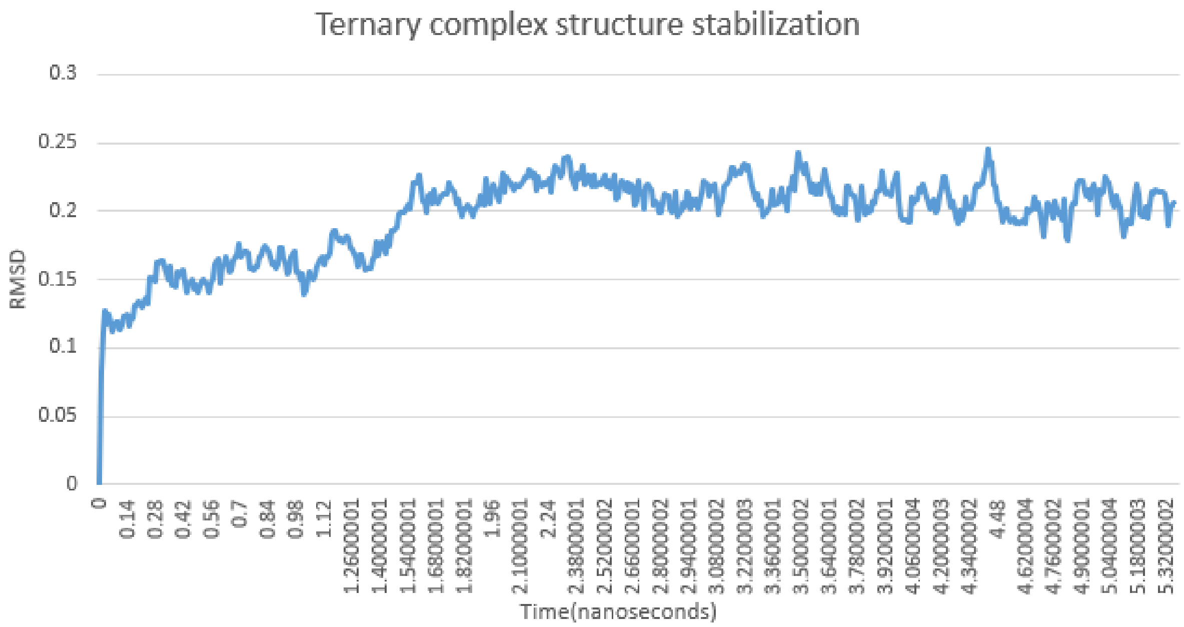

Figure 5 – Structure stabilization of ternary complex

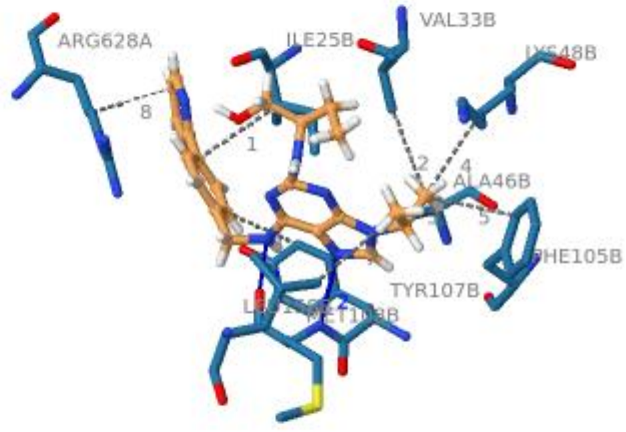

Figure 6 – RC8 interacting with DDB and CDK

Table 1 – RC8 interaction with DDB and CDK

| DDB (Chain_A) | Interaction Type | Interacting residues |
|---|---|---|
| | Hydrophobic | 928ARG |
| | Hydrogen bond | 950ASN |
| CDK (Chain_B) | Hydrophobic | 815TYR, 813PHE, 756LYS, 754ALA, 733ILE, 741VAL |
| | Hydrogen bond | 816MET |

Table 2 – Potential Molecular glue candidates for CDK degradation among purchasable drugs

| Drug Name | SMILES | Binding Energy |
|---|---|---|
| Tolvaptan | Cc1ccccc1C(Nc1ccc(C(N2CCC[C@H](c3cc(ccc23)[Cl])O)=O)c(C)c1)=O | -14.68 |
| Eltrombopag | CC1C(C(N(c2ccc(C)c(C)c2)N=1)=O)=NNc1cccc(c2cccc(c2)C(O)=O)c1O | -12.39 |
| Rivaroxaban | [H][C@]1(CNC(c2ccc(s2)[Cl])=O)CN(C(=O)O1)c1ccc(cc1)N1CCOCC1=O | -12.18 |
| Vilazodone | C(CCN1CCN(CC1)c1ccc2c(c1)cc(C(N)=O)o2)Cc1c[nH]c2ccc(C#N)cc12 | -12.07 |
| Lumefantrine | CCCCN(CCCC)CC(c1cc(cc2C(=Cc3ccc(cc3)[Cl])c3cc(ccc3c12)[Cl])[Cl])O | -11.78 |

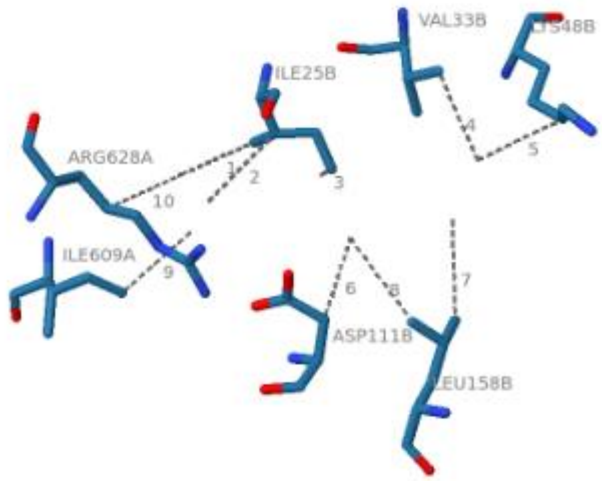

Figure 7 – Tolvaptan interacting with DDB and CDK

Table 3 - Tolvaptan interactions with DDB and CDK

| DDB (Chain_A) | Interaction Type | Interacting residues |
|---|---|---|
|  | Hydrophobic | ARG, ILE |
| CDK (Chain_B) | Hydrophobic | LYS, ASP, LEU, ILE, VAL |
| | Hydrogen bond | 816MET |

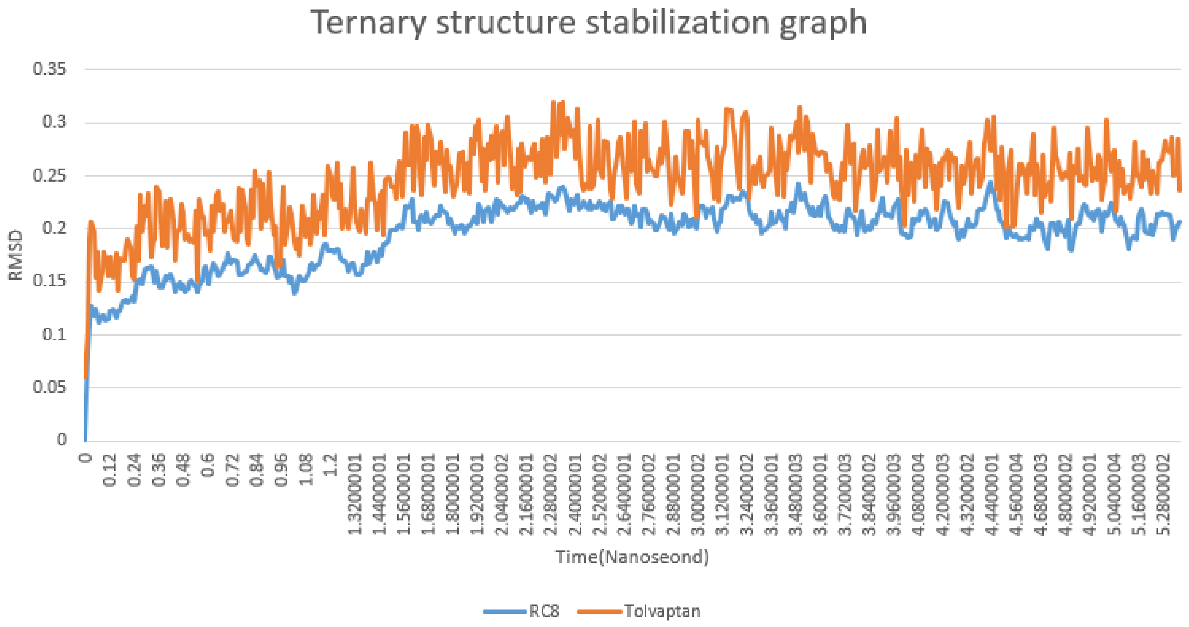

Figure 8. Ternary complex stabilization plot

Table 4 – Potential NCE Molecular glue candidates for CDK degradation

| 2D structure | Binding Energy | Carcinogenicity * | Drug induced liver injury* | Bioavailability* |
|---|---|---|---|---|
| 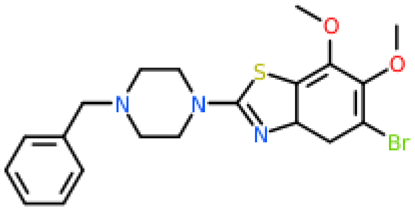 SAIT – MolGlue - 001 | -14.88 | 0.141 | 0.192 | 0.806 |
| 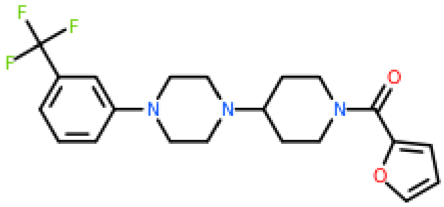 SAIT – MolGlue - 002 | -14.73 | 0.503 | 0.581 | 0.263 |
| 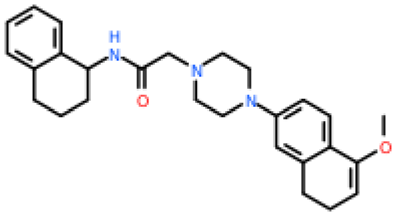 | -14.17 | 0.059 | 0.465 | 0.956 |

| | | | | |
|---|---|---|---|---|
| SAIT – MolGlue - 003 | | | | |
| 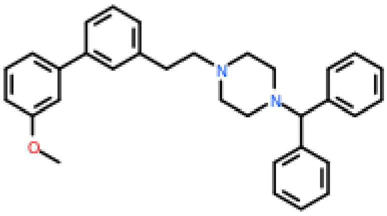 SAIT – MolGlue - 004 | -14.14 | 0.022 | 0.016 | 0.937 |
| 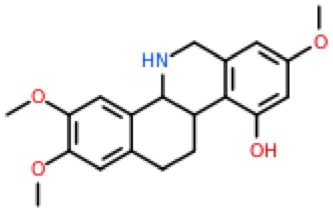 SAIT – MolGlue - 005 | -14.03 | 0.154 | 0.646 | 0.668 |

*These are probabilities in [0,1] range. Closer to 1.0 implies highly probable.

Table 5 - Tolvaptan interactions with DDB and CDK

| DDB (Chain_A) | Interaction Type | Interacting residues |
|---|---|---|
| | Hydrogen bond | 607ASN, 628ARG |
| CDK (Chain_B) | Hydrophobic | 48LYS, 105PHE, 158LEU, 168ALA |
| | Salt bridges | 111ASP |
| | Hydrogen bond | 109ASP |
| | Halogen bond | 35LYS |
